# Novel in situ seeding immunodetection assay uncovers neuronal-driven alpha-synuclein seeding in Parkinson’s disease

**DOI:** 10.1101/2025.04.05.647370

**Authors:** Maria Otero-Jimenez, Marcelina J Wojewska, Simona Jogaudaite, David Miller, Sandra Gray-Rodriguez, Grainne C Geoghegan, Laura Abelleira-Hervas, Tim J Viney, Barbara Sarkany, Djordje Gveric, Steve Gentleman, Javier Alegre-Abarrategui

**Author notes:** These authors contributed equally to this work (Marcelina J Wojewska, Simona Jogaudaite). Correspondence to: Javier Alegre-Abarrategui.

## Abstract

Aggregates of alpha-synuclein (α-syn) and tau propagate through template-induced misfolding in the brains of Parkinson’s (PD) and Alzheimer’s disease (AD) patients. Prion-like seeding is crucial in disease initiation and progression and represents a major target for drug-modifying therapies. The detection of aggregated α-syn and tau seeding activity with seeding amplification assays (SAAs) have remarkable diagnostic and research potential. However, current SAAs rely on bulk tissue homogenates or fluids, losing critical spatial and cellular resolution. Here, we report our novel *in situ* seeding immunodetection (*is*SID) assay that enables the visualization of α-syn and tau seeding activities with unprecedented morphological detail in intact biological samples. Using the *is*SID assay, we confirm seeding activity in the pathological aggregates of PD and AD, among others, while uncovering neuron-driven α-syn seeding events that precede the onset of clinical symptoms in PD. Our findings provide new fundamental insights into the pathogenesis underlying neurodegeneration and establish an invaluable tool for studying protein aggregation dynamics, with potential applications in biomarker discovery, diagnostics and drug testing.

## Introduction

Many common neurodegenerative disorders are characterized by the aggregation of misfolded proteins. These form pathological inclusions^1,2^, disrupting cellular function and eventually leading to neuronal cell death^2^.

Alpha-synucleinopathies are a group of neurodegenerative disorders characterized by the aggregation of alpha-synuclein (α-syn), a protein thought to play a physiological role in synaptic function^3^. These diseases include Parkinson’s Disease (PD), Parkinson’s Disease Dementia (PDD), Dementia with Lewy Bodies (DLB) and Multiple System Atrophy (MSA). The defining neuropathological feature of PD, PDD, and DLB, collectively termed “Lewy body disease” (LBD) is the presence of α-syn aggregates within neurons forming Lewy bodies (LBs) and Lewy neurites (LNs), which can be visualised histologically. However, glial pathology is becoming increasingly recognized. Recently, we have reported a distinct pattern of neuroanatomical progression, in which neurons are the first affected cell type, followed by astrocytes, when α-syn pathology becomes widespread^4^. In contrast, MSA is characterized by α-syn aggregating within oligodendrocytes, manifesting as glial cytoplasmic inclusions (GCIs) with a lower prevalence of neuronal cytoplasmic (NCIs) and nuclear inclusions (NNIs)^5,6^. Additionally, 8-17% of aged neurologically normal individuals exhibit LBs upon post-mortem examination, termed incidental Lewy body disease (iLBD), which is thought to represent a prodromal stage of PD^7^.

While the precise factors driving protein misfolding and aggregation remain incompletely understood, evidence suggests pathological aggregates of misfolded proteins (amyloids) can propagate in a prion-like manner^8^. This mechanism is hypothesized to be shared with other protein-misfolding diseases like tauopathies, which are characterized by the aggregation of hyperphosphorylated tau^9–11^. Prion-like seeding involves the induction of aberrant misfolding from the pathological aggregates to native proteins. This then leads to the formation of oligomeric species and fibrillar inclusions, facilitating disease initiation and progression^12^. However, whether all pathological forms present across cell types are equally seeding-competent.

The prion-like activity of α-synuclein, tau and other amyloids has been exploited to develop cell-based and cell-free seeding assays. Cell-based assays include using biosensor HEK293T cell lines which stably express α-syn or tau fused to cyan fluorescent protein (CFP) and yellow fluorescent protein (YFP). Liposome-mediated transduction of proteopathic seeds, either patient-derived or pre-formed fibrils (PFFs), into biosensor cells induces aggregation of CFP and YFP, generates fluorescence resonance energy transfer (FRET) signals as well as formation of cytoplasmic inclusions^13,14^. Cell-free seeding assays include protein misfolding cyclic amplification (PMCA) and real-time quaking-induced conversion (RT-QuIC), originally developed for the detection of misfolded prion proteins^15,16^. The PMCA assay involves cyclic sonication and incubation with vast excesses of normal brain homogenates containing endogenous monomers^15^. The RT-QuIC assay relies on a reaction buffer containing recombinant monomeric protein, the fluorescent dye thioflavin T (ThT), and samples containing the protein aggregates of interest^16–18^. Although originally developed using homogenized brain tissue, the assay has since evolved to be compatible with cerebrospinal fluid (CSF), skin, olfactory mucosa, and gut tissue^19–23^. The reaction mixture is subjected to orbital shaking and if pathological protein is present in the sample, it seeds the aggregation of the recombinant protein. The aggregation process involves cycles of elongation and fragmentation which are monitored in real time through the fluorescence emitted by ThT intercalating into the newly formed aggregates^24^. Since their development, cell-free seeding assays have been adapted for the detection of pathological prion-like proteins such as α-syn, tau, and TAR DNA-binding protein (TDP-43)^25,26^. Importantly, both the α-syn and tau RT-QuIC assays are able to successfully detect positive responses even in prodromal cases before the appearance of pathology and overt clinical symptoms^27–31^.

Despite its utility, the RT-QuIC assay requires bulk sample input and thus, spatiotemporal information about the seeding activity is lost. A detailed understanding of the cellular and subcellular distribution of seeding activity is critical in proteinopathies, as certain cells exhibit selective vulnerability^32–34^. For instance, more than a century after the discovery of LBs, it remains unknown if LBs and other types of α-syn inclusions possess seeding activity^35^. Recent studies have now demonstrated that α-syn seeding activity information can be obtained from formalin-fixed paraffin-embedded (FFPE) tissue, a widely used method to preserve and archive tissue^36–38^. Studies have confirmed seeding activity in brains fixed from only 18 hours to archival tissue fixed for over 20 years old^36–38^. However, the samples still required aggressive sample preparation steps, including tissue homogenization, sonication, deparaffinization, dissociation and heat-mediated antigen retrieval. Remarkably, these studies demonstrated that the seeding activity of FFPE tissue was comparable to that of frozen tissue and similarly minimal in control samples. Hence, as FFPE tissue sections are likely to preserve seeding activity, we next sought to develop a new assay to visualize seeding activity within intact tissue. For this, we developed the novel *in situ* seeding immunodetection assay (*is*SID), which leverages the endogenous seeding mechanism utilized in seeding amplification assays, combined with in situ visualization through immunostaining techniques. In this study, we describe the *is*SID assay for α-syn and tau, revealing for the first time the spatial and cellular distribution of seeding activity with fully preserved histomorphology in a cohort of PD, DLB, MSA, iLBD, and AD cases. Applying this novel assay, we have uncovered that neurons are presumably the main drivers of α-syn seeding although glial cells are also able to actively contribute to the seeding process. Additionally, we have successfully detected α-syn seeding in asymptomatic individuals with incidental α-syn pathology, demonstrating that α-syn seeding precedes the development of clinical symptomatology.

## Materials and methods

### *Post-mortem* human tissue

The cohort was composed of MSA (n = 12), DLB (n = 11), PD (n = 24), iLBD (n = 8), AD (n = 3), and healthy controls (n = 6). FFPE *post-mortem* human tissue was obtained from The Multiple Sclerosis and Parkinson’s UK Tissue Bank at Imperial College London and used under Research Ethics Committee Approval Ref. No 07/MRE09/72. Regions analyzed included the amygdala, midbrain, pons, medulla, cerebellum, hippocampus as well as superior frontal, anterior cingulate, temporal, and entorhinal cortices.

### *In situ* SID on tissue sections

This technique has a filed patent, application number GB2410222.0. For the *is*SID, FFPE-tissue sections were baked at 55°C for 45 minutes prior to dewaxing in xylene and rehydration in sequential concentrations of ethanol. The endogenous peroxidase activity was blocked at room temperature (RT). The reaction buffer was added to the tissue sections (containing his-tagged α-syn and ThT). The slides were placed into the plate holder, sealed with roofed chambers, and then incubated in a FLUOstar Omega microplate reader (BMG Labtech) under shaking or not shaking conditions. After the incubation, tissue sections were washed and blocked for 1 hour with 10% normal horse serum (NHS). The sections were then incubated with anti-6x-His tag antibody. Sections were visualized with ImmPRESS[R] HRP Horse Anti-Rabbit IgG Polymer Detection Kit, Peroxidase and ImmPact DAB Substrate HRP. Tissue sections were counterstained with Haematoxylin, dehydrated with ethanol and xylene prior to coverslipping with Epredia™ ClearVue™ Coverslipper (**Fig. 1**).

**Fig. 1:**
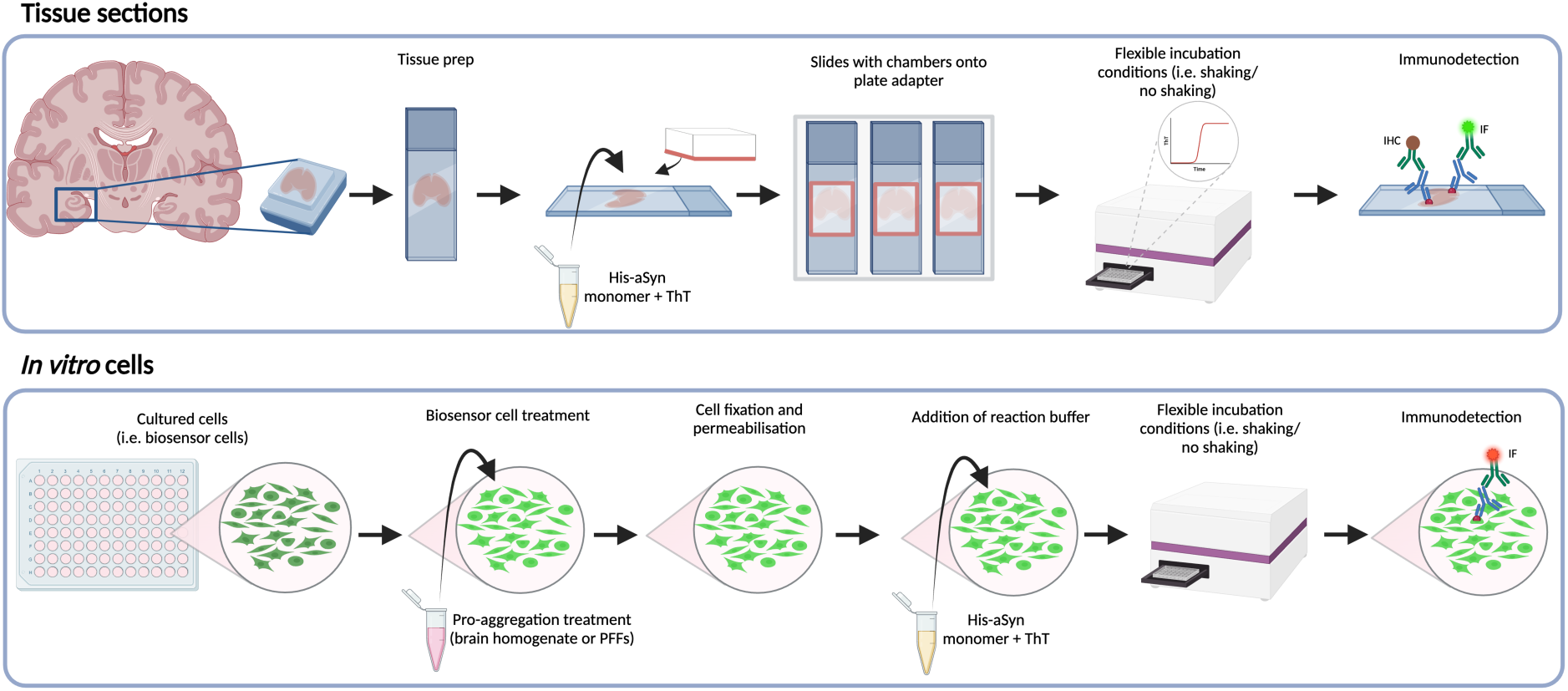
Workflow of the *is*SID assay on tissue sections and *in vitro* models. In tissue sections, slides undergo tissue preparation. Slides are placed into a plate adapter, sealed, and incubated with his-tagged monomeric substrate and ThT. The slides are incubated under flexible incubations (i.e. shaking or no shaking, in a plate reader. Following the incubation period, the his-tag is immunodetected by immunohistochemistry (IHC) or immunofluorescence (IF). *Is*SID can also be performed on cultured cells such as biosensor cells. For the biosensor cells, cells are treated with the pro-aggregation treatment (i.e. transduction complexes containing PFFs). After fixation with 4% PFA and permeabilization, cells are incubated with his-tagged monomeric substrate and the his-tag is immunodetected.

### *In situ* SID on cultured cells

HEK293T cell lines engineered to express α-syn (A53T) were kindly provided by Marc Diamond. For seed transduction, α-syn-expressing HEK293T cells were transduced with α-syn PFFs, opti-MEM and Lipofectamine 2000. After 24 hours, cells were fixed with 4% PFA and permeabilized for 30 minutes at RT. Following permeabilization, cells were incubated with his-tagged α-syn in a FLUOstar Omega microplate reader. Then, cells underwent blocking with 10% normal donkey serum (NDS). Cells were then incubated with primary antibody followed incubation with the fluorescent secondaries. Finally, wells were incubated with DAPI, and coverslips were mounted onto slides with VECTASHIELD® Antifade Mounting Medium (**Fig. 1**).

### Immunohistochemistry (IHC)

FFPE-tissue sections were dewaxed and rehydrated as previously described. The endogenous peroxidase activity was blocked in 0.3% H2O2 for 30 minutes at RT and antigen retrieval was performed in 100% formic acid for 10 minutes. The tissue sections were blocked in 10% NHS and incubated with primary antibody overnight at 4°C. Next day, the slides were incubated with ImmPRESS[R] HRP Horse Anti-Mouse IgG Polymer Detection Kit for 30 minutes at RT. ImmPact DAB Substrate HRP was added, and sections were counterstained with Haematoxylin, dehydrated with ethanol and xylene, before being coverslipped using Epredia™ ClearVue™ Coverslipper.

### Immunofluorescence

FFPE-tissue sections were baked prior to dewaxing in xylene and rehydration in ethanol. Antigen retrieval and the *is*SID assay were performed as previously described, omitting the ThT. Next day, after the roofed chambers were removed and the sections were blocked for 1 hour with 10% NDS. The sections were incubated with the antibodies for 4 hours at RT followed by incubation with the fluorescent secondaries for 2 hours. Slides were incubated with 0.1% Sudan Black before mounting with VECTASHIELD® Antifade Mounting Medium with DAPI.

### AS-PLA / tau-PLA

α-Syn or tau Proximity Ligation Assays (AS-PLA/tau-PLA)^39,40^ were performed as follows. Conjugates were generated by conjugating either anti-α-syn antibody MJFR1 (Abcam, ab209420) or anti-tau antibody TAU-5 (Abcam, ab80579) to PLA oligonucleotides using the Duolink® In Situ Probemaker PLUS (Sigma, DUO92009) and MINUS (Sigma, DUO92010) kits^39,40^. FFPE-tissue sections were deparaffinized and rehydrated as previously described. The endogenous peroxidase activity was blocked in 10% H_2_O_2_ at RT for 1 hour and followed by heat-mediated antigen retrieval in the microwave. Tissue sections were incubated with endogenous avidin and biotin blocking solution (Biolegend, 927301) as per manufacturer’s instructions before incubating with the provided PLA blocking solution for 1 hour at 37°C. Briefly, PLA conjugates were diluted in the probemaker diluent and incubated overnight at 4°C. Then, the assay was performed as described in Söderberg *et al.* (2006)^41^ with minor modifications. The next day, the sections were incubated with T4 ligase buffer, two connector oligonucleotide probes, adenosine triphosphate (ATP), and T4 ligase for 1 hour at 37°C. After washing with tris buffered saline (TBS) 0.05% Tween, rolling circle amplification was performed by adding the phi29 polymerase buffer, deoxynucleotide triphosphates (dNTPs), and phi29 polymerase then incubating for 2.5 hours at 37°C. Detection was then carried out by adding detection buffer containing SSC, polyadenylic acid (poly-A), bovine serum albumin (BSA), 0.5% Tween, and biotin-labeled oligonucleotide, followed by incubation for 1 hour at RT. Subsequently, the slides were incubated with Strepavidin-HRP solution (Abcam, AB64269), followed by development with Vector NovaRed substrate. and then counterstained, dehydrated, and coverslipped using the Epredia™ ClearVue™ Coverslipper.

### Image analysis

Brightfield images were acquired with a DS-Fi2 Digital Camera attached to a Nikon Eclipse 50i microscope, using a DS-U3 Digital Camera Controller and NIS-Elements image acquisition software. Immunofluorescence slides were imaged at 40x magnification with a numerical aperture of 0.95 on an Olympus BX63 scanning fluorescence microscope implementing the cellSens imaging software. Cells were imaged at 40x magnification with a numerical aperture of 1.25 on the Leica TCS SP8 confocal microscope implementing the Leica Application Suite X imaging software. To visualize FRET, the donor CFP fluorophore is excited at 458 nm, and its emission is collected with 480–520 nm filter. If the acceptor fluorophore YFP is in close proximity, energy transfer occurs, exciting YFP indirectly whose emission is then detected at 560 nm, confirming FRET.

### Quantification of cell-specific α-syn pathology

The classification of cell-specific α-syn pathology was performed like previously described^4^. In brief, stained tissue sections were scanned using the Leica Aperio AT2 slide scanner. QuPath software, version 0.4.4.^42^ and StarDist, version 0.3.2., a deep learning approach^43^, were used for the development of artificial neural network (ANN)-classifiers to assist in the supervised the detection of cell-type specific α-syn inclusions. Cells that were detected by StarDist and QuPath were used to train ANN-classifiers by manual annotation. After training of the ANN-classifiers, three ROIs corresponding to a 20x field of view were drawn, and the ANN-classifiers were applied.

### Statistical analysis

GraphPad Prism software was used for statistical analysis. The levels of α-syn pathology in neurons, astrocytes and oligodendrocytes detected by IHC and *is*SID were compared by two-way ANOVA was performed followed by Bonferroni’s *post-hoc* test. Statistical significance was set at p<0.05.

## Results

### *In situ* SID assay specifically localises endogenous α-syn and tau seeding activity in human brain tissue sections and cultured cells with intact morphological detail

To understand cell-type specific α-syn seeding activity in a spatial context, we developed the α-syn-*is*SID assay. We first explored if PD brain FFPE tissue sections had seeding-competent α-syn that could be visualized *in situ.* For this, tissue sections were subjected to a modified RT-QuIC protocol where slides were incubated with his-α-syn as a substrate (**Fig. 1**). Cases with α-syn pathology, as determined by IHC and AS-PLA, exhibited an increase of ThT signal in comparison to cases without pathology or in which his-α-syn monomers were omitted (**Fig. 2a**).

**Fig. 2:**
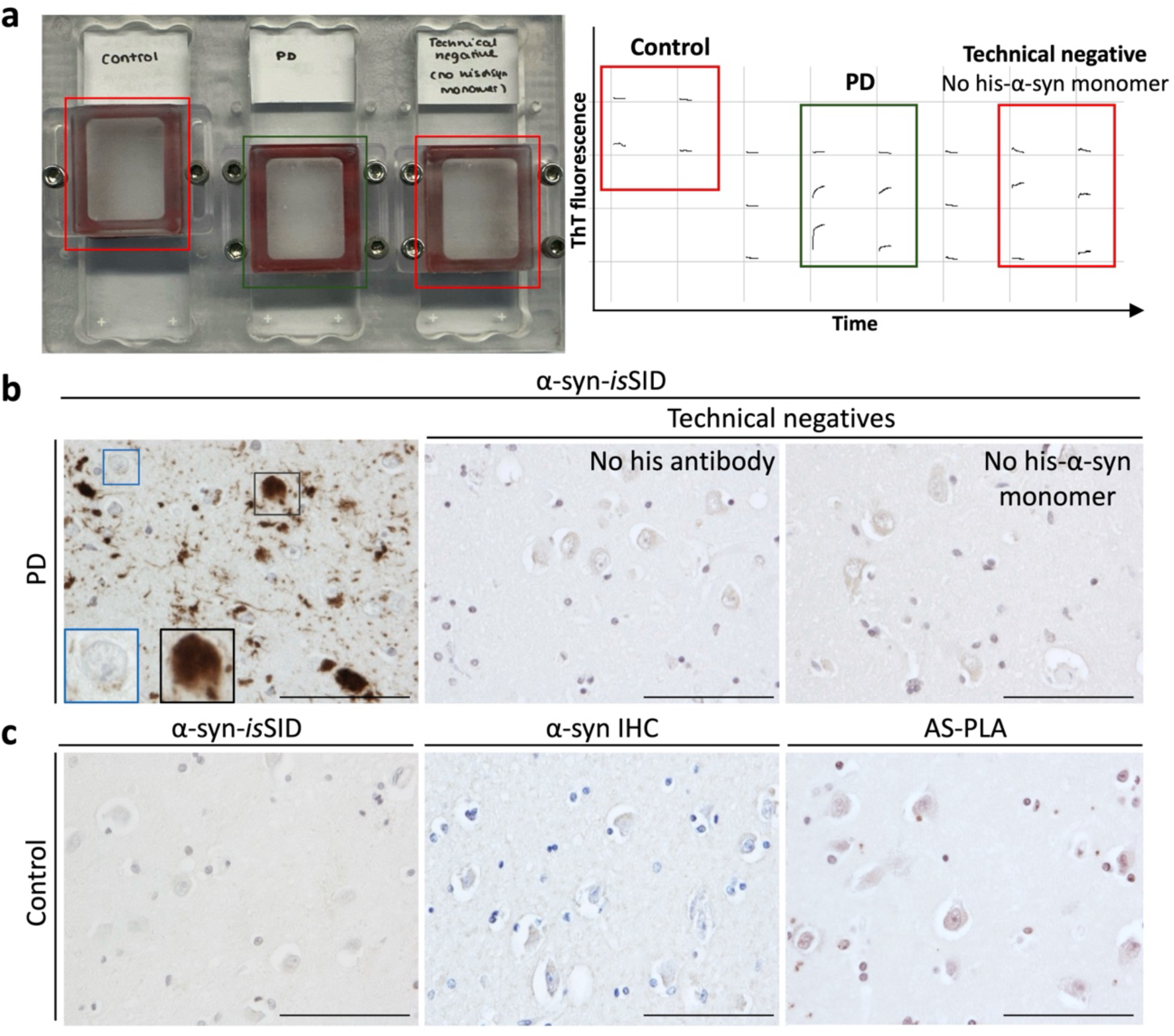
Validation of the α-syn-*is*SID specificity. Overview of the α-syn-*is*SID assay set up (left). Marked increase in ThT fluorescence signal was exclusively detected in cases with previously characterized pathology and a representative PD case shown (green box). No relative increase in ThT signal is seen in the representative control case or in the technical negative where no his-α-syn monomer was added, (red boxes) (**a**). In the amygdala of a PD case, extensive seeding-competent α-syn pathology was observed (grey square), while neurons lacking α-syn pathology (blue square) were also present confirming signal specificity. Omission of the anti-his antibody or recombinant his-α-syn substrate during incubation abolished signal, confirming that the assay specifically detects newly aggregated monomeric his-α-syn *in situ* (**b**). Control cases without α-syn pathology, as confirmed by α-syn IHC and AS-PLA, showed no signal in α-syn-*is*SID (**c**). Scale bar is 100 µm and insets represent magnified areas.

We attempted to visualize nascent α-syn aggregates seeded *in situ* using IHC with antibodies against the his tag. However, slides assayed in this way were poorly secured in the plate reader and inconsistently retained the assay buffer. In addition, we observed α-syn-*is*SID signal appearing within tissue regions located outside, and particularly the underside, of the cylinder. To maintain consistent humidity during slide incubation, whether inside or outside the plate reader, and to also maximize the assay area, we developed prototypes of a slide holder, roofed chamber, silicone foam gasket and cylinders with thicker walls.

The prototypes allowed consistent visualization of newly formed α-syn aggregates seeded *in situ* (**Fig. 2a**). Removal of the his-α-syn monomeric substrate resulted in no signal, indicating that the assay’s signal is reliant on the *de novo* aggregation of monomeric his-α-syn seeded *in situ* by endogenous α-syn aggregates. Removal of the anti-his antibody resulted in no signal, indicating that the assay exclusively detects newly aggregated monomeric his-α-syn (**Fig. 2b**). The α-syn-*is*SID signal was solely observed in the presence of pathological α-syn. No signal developed in control cases lacking α-syn pathology, as assessed by α-syn-IHC and AS-PLA (**Fig. 2c**).

To further validate the *is*SID assay for detecting α-syn seeding activity and to establish if we could visualize seeding activity *in situ* in a cell culture model, we applied α-syn-*is*SID to α-syn biosensor cell lines^13,14^. After treating the α-syn biosensor cells with PFFs and performing *is*SID, we observed cytoplasmic inclusion formation and FRET signal, indicating successful seeding (**Fig. 3**). Following cell fixation and permeabilization, we performed α-syn-*is*SID with his-α-syn followed by immunofluorescence using the anti-his antibody (**Fig. 1**). Strikingly, the *is*SID signal colocalized with the FRET signal and the induced inclusions (**Fig. 3**). These findings confirm that α-syn-*is*SID effectively detects seeding activity in both tissue samples and *in vitro* cell culture models. Importantly, no signal was observed in cells lacking his-tagged α-syn monomers, further demonstrating the assay’s specificity (**Fig. 3**).

**Fig. 3:**
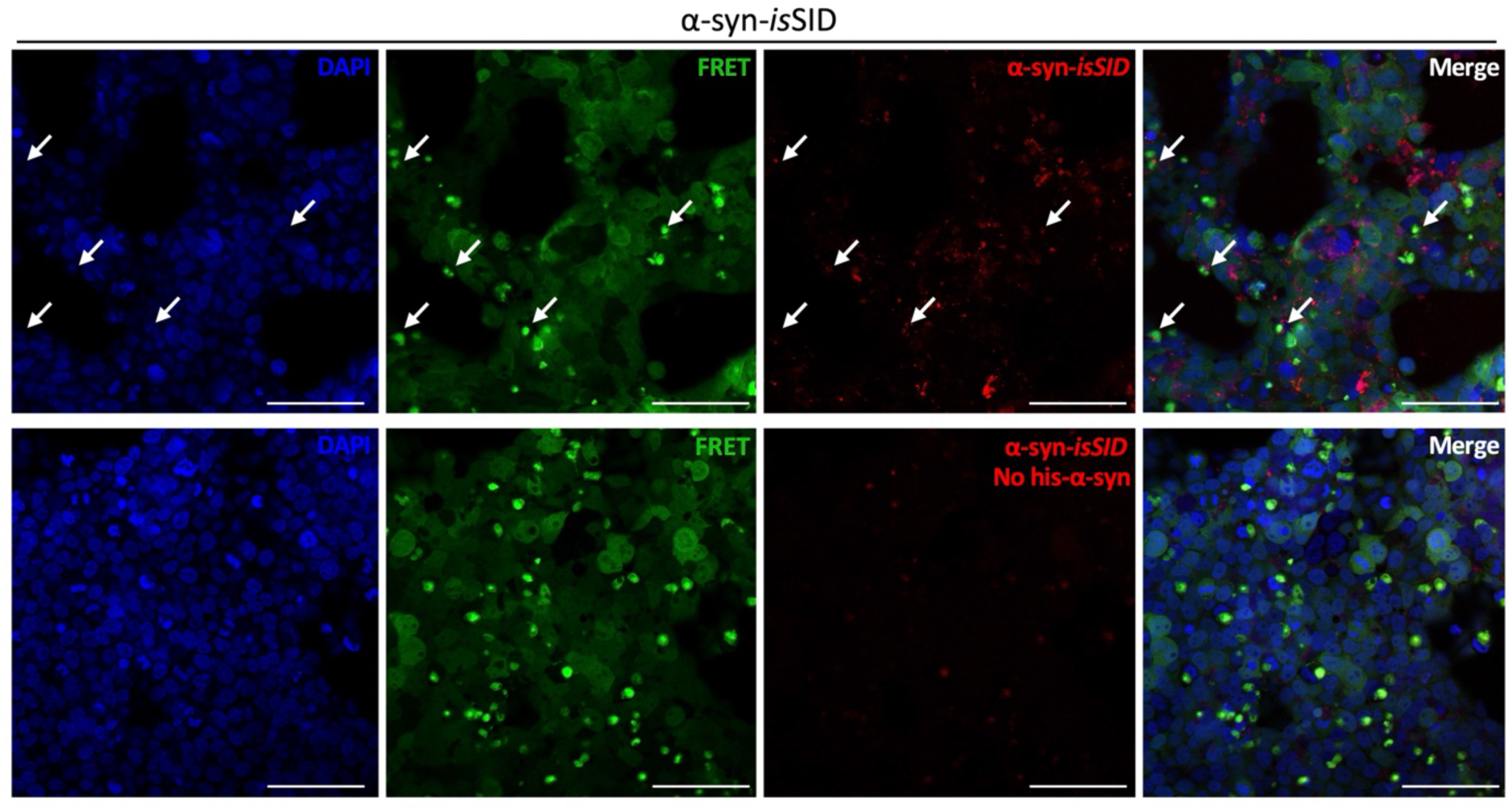
Validation of α-syn-*is*SID in cultured cells using FRET imaging. HEK293T α-syn biosensor cells were treated with α-syn PFFs and subjected to isSID. FRET was observed by exciting CFP at 405 nm and detecting YFP emission at 525–575 nm, indicating energy transfer seen as green fluorescence inclusions. Pathological inclusions in α-syn biosensor cells displayed a positive α-syn-*is*SID signal (indicated in red), which was absent when his-α-syn monomer was not added. Scale bar is 100 µm.

We then wanted to investigate if α-syn-*is*SID could be adapted to investigate seeding activity using other α-syn protein substrates such as the A53T mutant^2,44^. The A53T α-syn mutant protein was successfully and effectively shown to be used to explore α-syn seeding *in situ*, revealing *is*SID signal in LBs, astrocytes, and oligodendrocytes (**Supplementary Fig. 1**).

While the main purpose of this study was to establish the spatial distribution of endogenous α-syn seeding activity, we wanted to test whether this novel assay could be extended to other proteins with prion-like seeding properties. By modifying the *is*SID protocol to incorporate the use of his-tagged tau monomers as the substrate, we applied the assay to a cohort of AD cases. The tau-*is*SID assay successfully demonstrated that tau seeding activity can be visualized *in situ* (**Fig. 4a**). Similarly to α-syn, tau-*is*SID did not produce any signal in the control conditions lacking either the his-tagged monomeric protein or anti-his antibody, confirming the assay’s specificity and excluding non-specific background binding. No detectable tau seeding activity was observed in cases with no tau pathology as defined by both IHC and tau-PLA (**Fig. 4b**).

**Fig. 4:**
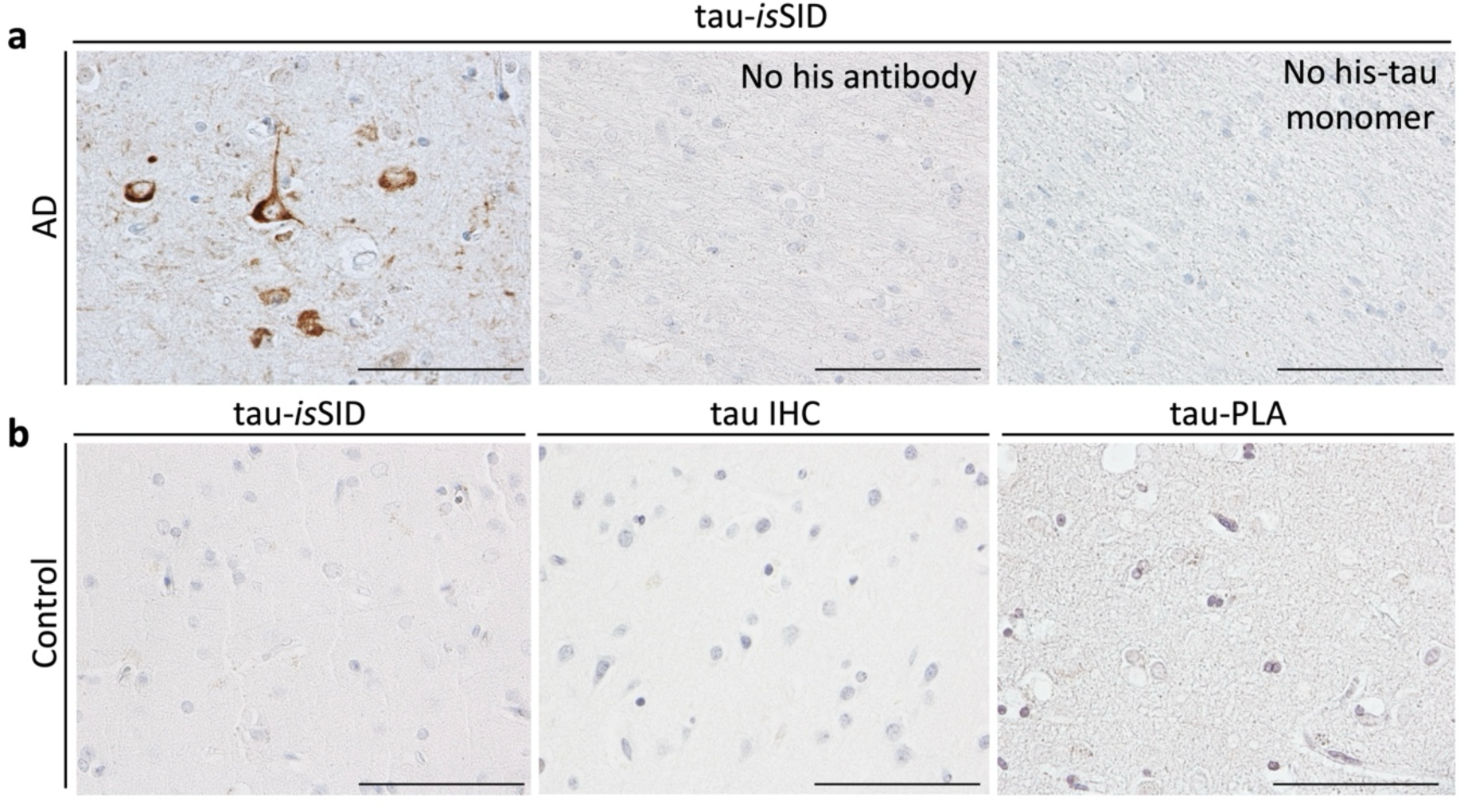
Validation of the tau-*is*SID specificity. Tau-*is*SID signal was observed in the temporal cortex of an AD case with previously characterized tau pathology, confirming signal specificity. Omission of either the anti-his antibody or monomeric his-tau substrate during incubation resulted in loss of signal, demonstrating that the assay signal specifically detects newly aggregated recombinant his-tau *in situ* (**a**). Control cases without detectable tau pathology, as confirmed by either by tau IHC and tau-PLA, displayed no tau-*is*SID signal (**b**). Scale bar is 100 µm.

Shaking is a key factor required for bulk SAAs. Therefore, we sought to explore whether shaking was essential in our α-syn and tau *is*SID assays. We observed that α-syn-*is*SID was able to detect α-syn seeding activity *in situ* under non-shaking conditions, with results comparable to those obtained using the shaking protocol. In contrast, the tau-*is*SID assay produced significantly weaker signals under non-shaking conditions, and only dot-like foci of seeding activity in the soma of neurons and in the neuropil remained detectable (**Supplementary Fig. 2**). All experiments described in this manuscript, except where indicated, were performed under shaking conditions.

### *In situ* SID assay reveals both neuronal and glia α-syn inclusions, and tau aggregates possess prion-like seeding activity

After validating the specificity of the α-syn-*is*SID and tau-*is*SID assays, we aimed to address some unresolved questions including the seeding capacity of LBs and other neuronal pathologies as well as whether glial cells harbored seeding-competent aggregates. By applying α-syn-*is*SID to PD and DLB cases, we observed seeding-competent α-syn pathological inclusions present in several cell types. Importantly, using this novel technique, we were able to confirm that LBs are not inert. We also detected seeding-competent α-syn neuronal structures in the shape of LNs. In addition to neuronal α-syn pathology, α-syn-*is*SID revealed that α-syn glial inclusions, including astrocytes and oligodendrocytes, have seeding capacity (**Fig. 5**). The morphologies of seeding-competent astrocytic α-syn inclusions included all our recently described novel morphologies^4^. We confirmed by double labelling immunofluorescence of α-syn-*is*SID with a variety of cell markers in PD cases, that α-syn-*is*SID signal was present in neurons, astrocytes, oligodendrocytes, as well as in microglia (**Supplementary Fig. 3**). Furthermore, we detected α-syn dot-like foci of seeding activity scattered in the neuropil and in the perikaryal cytoplasm of some neurons, which may represent early maturation stages in the process of LN and LB formation, respectively111^45^1 (^45^neuropil dot-like seeding foci predominately co-localized with neurofilament and synaptic markers, indicating their neuronal, and more specifically, synaptic, location (**Supplementary Fig. 4**). Moreover, we observed positive signal surrounding neuritic amyloid beta (Aβ) plaques^46,47^.

**Fig. 5:**
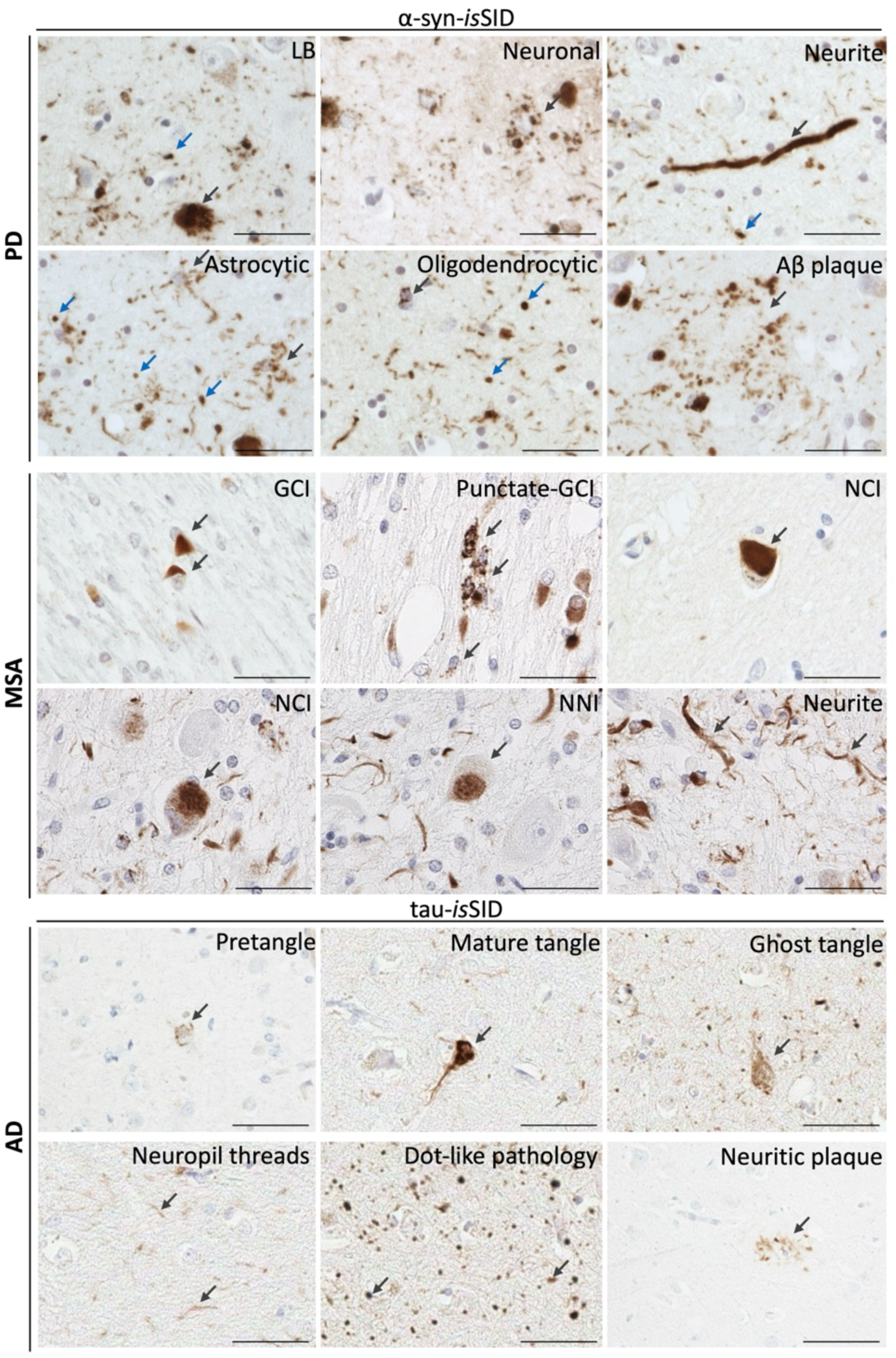
Is SID confirms that pathological α-syn and tau inclusions possess seeding potential. In PD cases, α-syn-*is*SID labelled pathological α-syn aggregates in LBs, neuronal, and neuritic inclusions, as well as in astrocytes, oligodendrocytes and on the periphery associated with an Aβ plaque. Blue arrows indicate dot-like pathology. In MSA cases, the α-syn-*is*SID confirmed seeding capacity in GCIs, punctate-GCIs, as well as neuronal inclusions including NCIs, NNIs and neurites. In AD cases tau-*is*SID detects seeding potential within pathological tau structures, including pretangles, mature neurofibrillary tangles, ghost tangles, neuropil threads, dot-like pathology, and neuritic plaques. Scale bar is 50 µm.

The α-syn-*is*SID was next applied to investigate whether α-syn aggregates in MSA had seeding activity. We revealed that α-syn aggregates present in oligodendrocytes, in the form of GCIs, had seeding capacity. Interestingly, we observed a population of oligodendrocytes displaying punctate cytoplasmic seeding activity (“punctate GCIs”). Neuronal inclusions were also composed of seeding-competent α-syn, including NCIs and NNIs, alongside neurite-like pathology. This confirms that despite GCIs representing the dominant pathology in MSA, seeding potential is not limited to oligodendrocyte inclusions but also to the less prevalent neuronal α-syn pathology (**Fig. 5**).

While alpha-synucleinopathies were the focus of this study, we sought to investigate if our findings with α-syn-*is*SID had a parallel in tau pathology. In AD cases, the tau-*is*SID assay detected seeding-competent tau pathology in pretangles, mature neurofibrillary tangles, ghost tangles, neuropil threads, and the dystrophic neurites of neuritic plaques. In addition, and similarly to α-syn, dot-like tau seeding activity foci were observed scattered in the neuropil and in the perikaryal cytoplasm of some neurons (**Fig. 5**). We next stained consecutive sections of PD and AD brains and labelled with α-syn/tau-*is*SID, AS/tau-PLA and α-syn/tau-IHC, respectively. Pathological aggregates of α-syn and tau (LBs and NFTs) were detected by their respective *is*SID assays, as well as by PLA and IHC. However, both the α-syn and tau dot-like neuropil foci of seeding activity observed with *is*SID appeared to correspond more readily with pathology detected by PLA than by IHC (**Supplementary Fig. 5**).

### Neuroanatomical and cell type distribution of α-syn seeding activity at different stages of PD and in MSA

We next investigated the neuroanatomical and cell type distribution of α-syn seeding activity present in PD across various disease stages, and in MSA. Regions affected at different stages of pathogenesis were captured, namely the medulla, pons, midbrain (SN), cerebellum, amygdala, cingulate and entorhinal cortices and the hippocampus.

In PD, brainstem-type LBs exhibited seeding capacity across the medulla, locus coeruleus (LC) and the SN. Notably, the halo of nigral LBs seen in dopaminergic neurons was labeled. Additionally, glial α-syn inclusions and α-syn-positive fibre tracts were detected in the medulla. The amygdala was a locus for the presence of various cellular α-syn inclusions displaying seeding-competence across LBs, neuronal inclusions, LNs, as well as astrocytes and oligodendrocytes. Similarly, cortical LBs demonstrated seeding competence alongside glial inclusions in the cingulate and entorhinal cortices. In the hippocampus, seeding activity was observed throughout, with the neuronal populations and LNs present in the CA2/3 region and the entorhinal cortex displaying the most severe involvement. Lastly, in the cerebellar white matter, a region largely unaffected in PD, the α-syn-*is*SID revealed a “dot-like” labeling in the neuropil (**Fig. 6**).

**Fig. 6:**
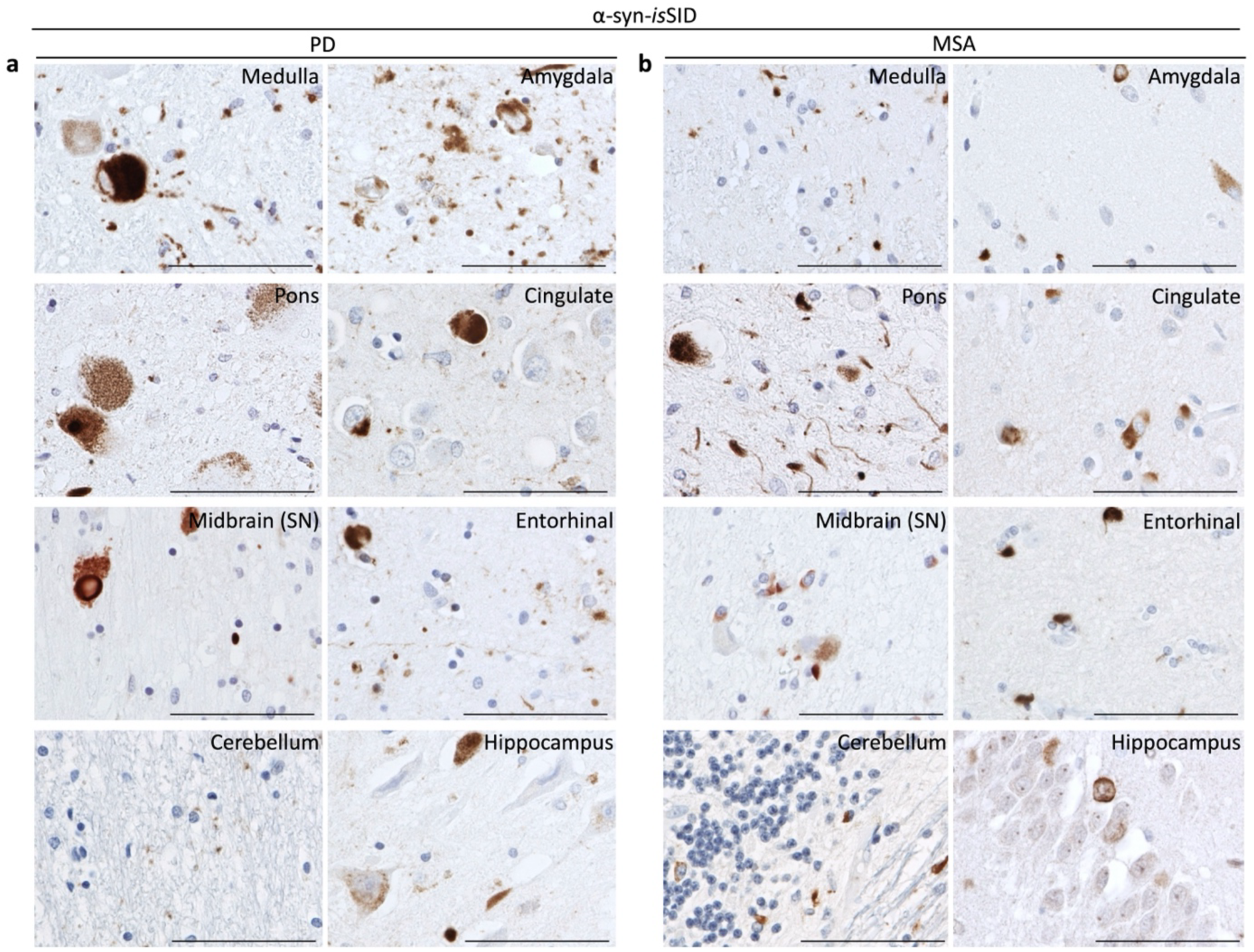
Distribution of α-syn seeding activity across distinct neuroanatomical brain regions in PD and MSA cases using the α-syn-isSID assay. Representative image illustrates the seeding activity assessed in multiple brain regions including the medulla, pons, midbrain (SN), cerebellum, amygdala, cingulate cortex, entorhinal cortex, and the hippocampus using α-syn-isSID assay in PD (**a**) and MSA cases (**b**). Scale bar is 100µm.

In MSA, we detected seeding-competent GCIs with diverse morphologies across all examined brain regions. Neuronal α-syn inclusions in the form of NNIs and NCIs alongside GCIs were found in the medulla to display α-syn seeding activity. Extensive α-syn-*is*SID signal was present in the pons, including neuronal inclusions and neuritic pathology throughout all the transverse fiber tracts in the base of pons. Seeding-competent neuronal inclusions were present in the amygdala and in cortex, in addition to the ring-like neuronal inclusions in the dentate gyrus. In the cerebellum, α-syn seeding activity was present in GCIs, “punctate GCIs”, as well as in the neuronal α-syn inclusions (**Fig. 6**).

### *In situ* SID assay reveals increased α-syn seeding activity in neurons as compared to glia in PD

Next, we aimed to investigate the relative proportion of neurons, astrocytes, and oligodendrocytes possessing seeding-competent α-syn pathology. For this, we compared the number of these cell types possessing α-syn seeding activity, as detected by α-syn-*is*SID, to the number of each cell type harboring IHC-positive α-syn inclusions. To correlate between these measures, we used consecutive tissue sections from the amygdala of both PD and MSA cases.

In PD cases, significantly more neurons displayed α-syn seeding compared to those harboring IHC-positive α-syn inclusions. No significant differences in the number of astrocytic and oligodendrocytic α-syn inclusions were detected between the techniques. In MSA cases, α-syn-*is*SID detected similar levels of oligodendrocytic and neuronal α-syn inclusions to IHC. Astrocytic α-syn inclusions were negligible with both techniques in MSA cases (**Fig. 7a, b**).

**Fig. 7:**
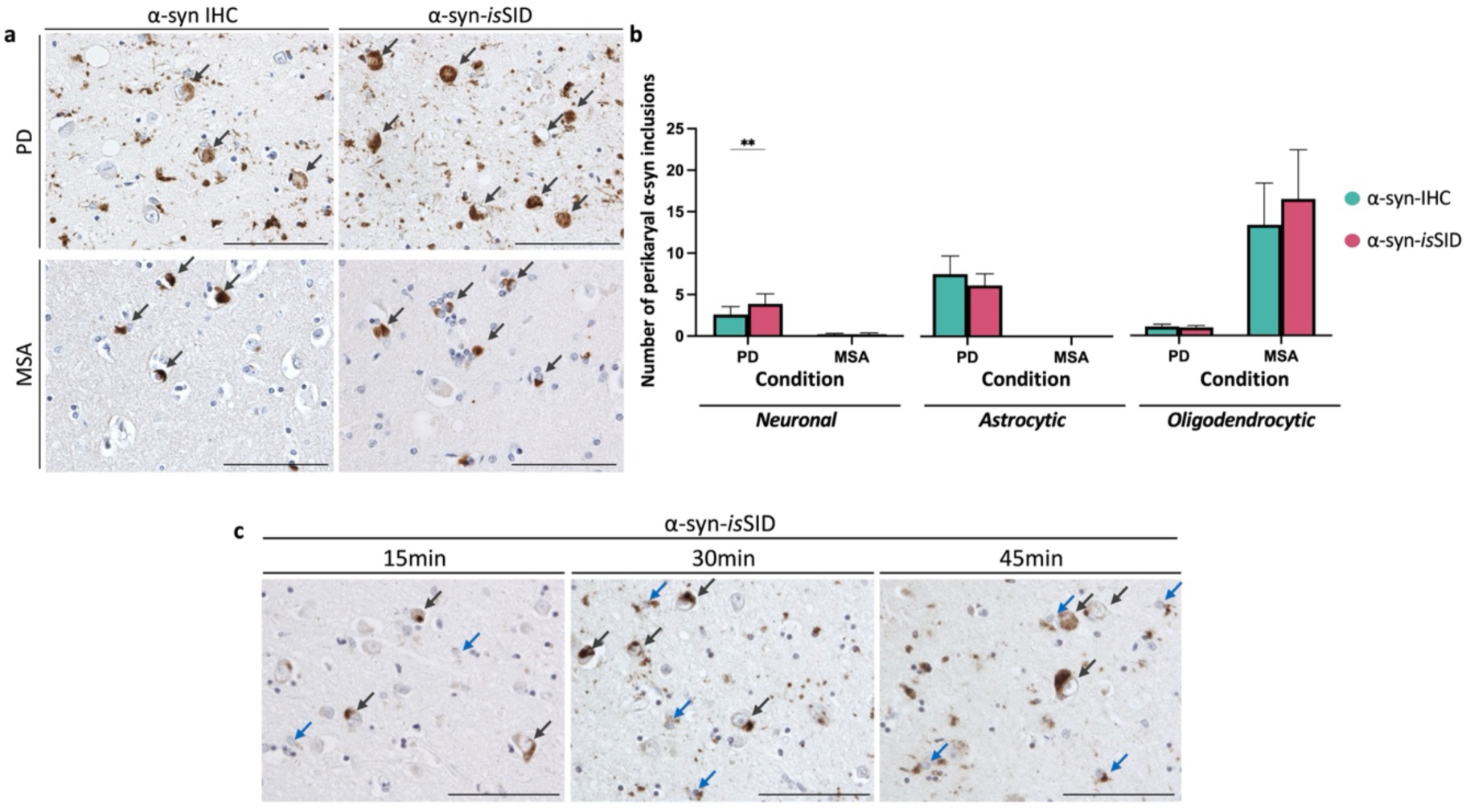
Neuronal α-syn pathology exhibits higher seeding activity than glial pathology in PD. α-syn aggregates and their seeding capacity were analyzed in the amygdala of PD and MSA cases using α-syn IHC and α-syn *is*SID. Representative images of neuronal inclusions detected by α-syn-isSID in PD and MSA (grey arrows) (**a**). Quantification of α-syn-isSID inclusions revealed more neuronal pathology in PD compared to IHC, whereas in MSA, both techniques detected similar levels of α-syn pathology. Oligodendrocytic and astrocytic α-syn pathology showed no significant differences between IHC and α-syn-isSID in either PD or MSA cases. Statistical analysis was performed using two-way ANOVA followed by Bonferroni post-hoc test (* p<0.05; ** p<0.01). Bars represent mean + standard error of the mean (**b**). Consecutive PD sections subjected to short α-syn-isSID incubation times displayed more prominent neuronal α-syn inclusions at 15 minutes compared to glial inclusions, although they were still detectable and increased over time. Grey arrows indicate neuronal α-syn inclusions; blue arrows indicate glial α-syn inclusions (**c**). Scale bar is 100 µm.

Both neurons and glial cells displayed α-syn seeding capacity, although additional α-syn seeding-competent neurons were revealed by α-syn-*is*SID in PD cases. To further determine any differential α-syn seeding activity of neuronal and glial α-syn inclusions, α-syn-*is*SID was performed with a time series of incubation periods (15 minutes, 30 minutes, and 45 minutes) in PD cases. These short incubation periods were selected to reveal any differential order of labeling of inclusion types. With 15 minutes of incubation, we observed incipient but widespread α-syn-*is*SID signal predominantly in neurons, although levels were reduced compared to the standard incubation protocol. Moreover, labeling of glia α-syn pathology was significantly more focal and weaker than neuronal pathology. However, the number of glial inclusions, as well as overall burden labeled by α-syn-*is*SID, was detected more readily with increasing incubation times of 30 minutes and 45 minutes. These results suggest neuronal α-syn pathology has increased seeding activity than glial cells (**Fig. 7c**).

### Seeding-competent α-syn is found in asymptomatic cases with incidental α-syn pathology

In order to assess the cell type distribution of α-syn seeding activity at early PD disease stages in the brain, we examined the brainstem of cases with Braak stage 1-3 pathology using the α-syn-*is*SID assay. These cases represented asymptomatic individuals with pathology restricted to the brainstem and categorised as incidental LB disease (iLBD) cases.

In the medulla, pons (LC), and the SN of these incidental LB-pathology cases, and in line with their Braak stage, α-syn seeding activity was detected in neuronal α-syn pathology (**Fig. 8a**). Oligodendrocytic α-syn seeding was scarce and no observable α-syn seeding activity was seen in astrocytes. α-Syn-*is*SID signal was present in neuromelanin-containing dopaminergic neurons, indicating seeding capacity (**Fig. 8a**). Additionally, we observed several patterns of α-syn-*is*SID signal in dopaminergic neurons, ranging from punctate pattern to a more condensed structure. We propose that these morphologies, which parallel the morphogenesis proposed for LBs^48,49^, likely represent the temporal evolution of α-syn seeding activity in dopaminergic neurons. α-Syn seeding activity would therefore start as a few punctate foci dispersed in the cytoplasm. Foci of seeding activity progressively became more abundant and larger, filling most of the cytoplasm, finally coalescing into LBs (**Fig. 8b**).

**Fig. 8:**
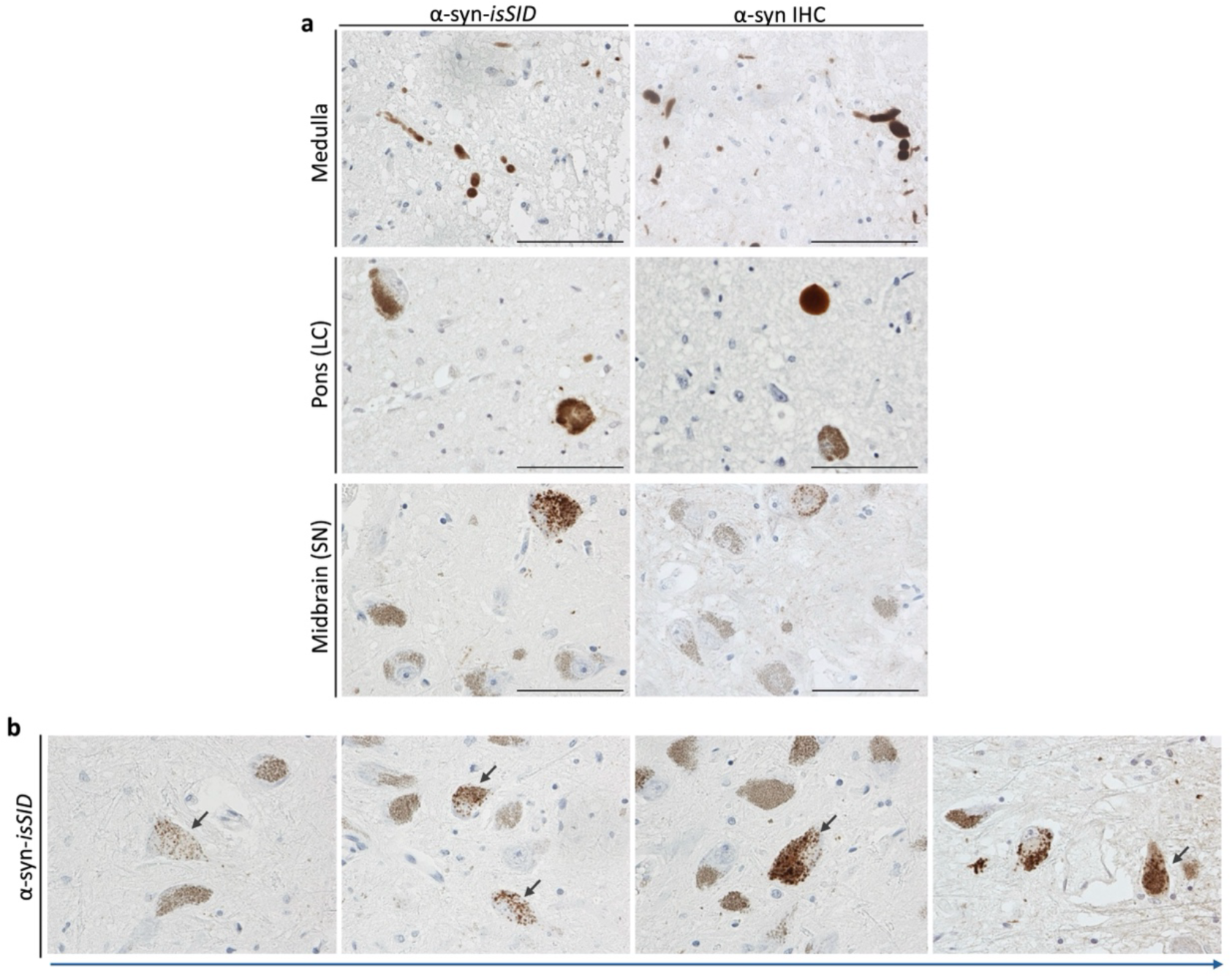
α-Syn seeding activity is detected in the brainstem of incidental cases with LB-pathology. Incidental cases with LB-pathology but no clinical symptomatology had seeding-competent α-syn pathology as shown by α-syn-isSID. Pathology detected by α-syn IHC is also shown (**a**). Proposed model of seeding and maturation stage of α-syn seeding in neuromelanin-positive neurons. α-Syn seeding is morphologically observed first in a punctate pattern, which progressively increments in size and number until covering the whole neuronal cytoplasm and eventually forming LBs (**b**). Scale bar is 100 µm.

## Discussion

Neurodegenerative proteinopathies are characterized by the ability of misfolded aggregated proteins to seed the conversion of endogenous proteins into their pathological counterparts. However, current methodologies that detect the seeding activity of biological samples lack spatial and cytoarchitectural resolution. Here, we describe our novel assay *is*SID for α-syn and tau proteins, which enables the direct visualization *in situ* of α-syn and tau seeding activities, resolving its spatial and cellular distribution.

Growing evidence indicates that the process of protein aggregation and seeding is central to the pathogenesis of neurodegenerative proteinopathies. However, the biological seeding activity of distinct cell types and inclusion subtypes has remained elusive, particularly regarding the ongoing debate over whether LBs exert a toxic or defensive role^50,51^. Here, we provide direct visual confirmation that the neuropathological hallmarks of PD/DLB, MSA and AD such as LBs, GCIs and NFTs, respectively, have biologically active seeding capacity. Additionally, we demonstrate that less common types of pathology such as glial inclusions in PD as well as neuronal inclusions (NCIs and NNIs) in MSA, also exhibit seeding activity. Specifically, in PD, our findings confirm that LBs are neither inert nor dormant structures, and despite their compact, insoluble structure they can actively promote pathological protein aggregation. The observation of neurons completely devoid of seeding competent α-syn next to neurons heavily burdened with pathological α-syn suggests that propagation likely follows defined anatomical routes rather than occurring via passive diffusion^52–54^. Small seeding activity foci were also seen in the cytoplasm of a proportion of neurons and oligodendrocytes in PD and MSA, respectively. These neuronal foci and previously unreported “punctate-like GCIs” presumably represent early stages preceding LB formation and early-stage GCIs^45^. However, we cannot ascertain what proportion of neurons and oligodendrocytes harbouring these punctate seeding foci will mature into fully developed inclusions or undergo cell death^45^. Increasingly confluent small foci of α-syn seeding activity were particularly observed in the soma of a proportion of monoaminergic neurons in the SN and in the LC, which supports the proposed cellular maturation model. Given that monoaminergic neurons in the SN and in the LC are among the most selectively vulnerable neuronal populations, these neurons might have intrinsic properties predisposing them to α-syn seeding, even at early disease stages, ultimately contributing to their progressive dysfunction and death. In addition, we identified dot-like foci of foci of α-syn seeding activity scattered in the neuropil which partially colocalize with presynaptic and neurofilament markers, suggesting at least a proportion of them reside in neuronal projections and synapses^55–57^. This may relate to the synaptic location of toxic oligomers and potential transsynaptic spread of aggregates. Moreover, α-syn-*is*SID signal was observed surrounding neuritic plaques with a morphology indicative of an association with dystrophic neurites^46,47^. Notably, the non-amyloid beta component of AD amyloid (NAC) region of α-syn has been reported as an integral component of Aβ plaques^55^. The observed seeding activity of α-syn alone may be sufficient to seed full^58^α-syn, in line with *in vitro* findings. Nevertheless, the possibility of α-syn and Aβ cross-seeding cannot be completely excluded.

*In vitro* studies have demonstrated that glia can actively propagate pathology; however, we now provide the first direct evidence that endogenous glial inclusions possess seeding capacity. These findings challenge the entirely neuron-centric view of PD by highlighting glia as active contributors to disease progression rather than passive bystanders. Recently, we have described different morphologies of astrocytic α-syn pathology in PD which correlated with the density of astrocytic α-syn pathology and here we show all six morphologies of astrocytic pathology exhibit seeding activity. Furthermore, our observation of α-syn-*is*SID signal present in microglia could support their involvement in disease dissemination,^59^ but, whether seeding-competent glia are actively promoting pathology or if they sequester toxic α-syn aggregates as a neuroprotective measure remains to be understood. The time required to reach the detection threshold during the lag phase is commonly used as a surrogate measure of seeding activity and seed concentration^60,61^. Hence, to investigate differential seeding activity between neuronal and glial α-syn pathology, we conducted the α-syn-*is*SID assay using very short incubation times. At the earliest timepoints, we observed a higher prevalence of neuronal compared to glial inclusions, suggesting a higher seed concentration within neuronal pathology or that neuronal-derived seeds have higher activity. Additionally, we detected more α-syn seeding-competent neurons than neuronal perikaryal lesions detectable by IHC. These results suggests that morphologically intact neurons can harbour active seeding foci not detectable by α-syn IHC, perhaps due to their small size. It could be speculated that these early seeding-competent forms may relate to the early aggregates we previously reported using AS-PLA^39^. iLBD is considered a pathological precursor of PD^62^ and our detection of α-syn seeding in iLBD cases confirms that seeding is already present at the presymptomatic stages^63^. This aligns with studies demonstrating that prodromal cases, such as RBD and PAF, are able to seed in SAAs despite lacking overt pathology^64–66^. Furthermore, our finding that neuronal α-syn seeding vastly predominated in iLBD cases, with very rare α-syn-*is*SID positive oligodendrocytic α-syn inclusions and no detectable astrocytic α-syn seeding, further support neurons being the main drivers of α-syn seeding. We recently reported that neuronal α-syn pathology appears first, followed closely by oligodendrocytic α-syn pathology, whereas, astrocytic α-syn pathology emerges once neuronal and oligodendrocytic pathology is established. In this study, we demonstrate that this neuroanatomical pattern of progression is also reflected at the seeding level, reinforcing the concept that α-syn spread follows a defined, hierarchical cellular trajectory.

While in this study we mostly focused on α-syn, we hypothesised that our method could be applied to other proteinopathies such as tau. In addition, we wondered if the endogenous neuronal and glial α-syn seeding revealed in PD and MSA cases was parallel to tau seeding activity in AD. Using tau-*is*SID we found, in contrast to α-syn, that tau seeding activity in AD seemed to be mostly restricted to neuronal elements, such as mature neurofibrillary tangles and neuropil threads, as well as prominent dot-like seeding activity foci scattered in the neuropil. However, tau seeding activity needs to be further explored in a larger cohort, including AD and other tauopathies such as Progressive Supranuclear Palsy characterized by tau pathology in glia, and differential neuroanatomical involvement.

Neurodegenerative diseases are becoming increasingly prevalent, with the key challenges including the lack of early diagnostic methods and effective treatments. Early diagnosis is crucial for the identification of novel biomarkers, the development of new therapeutics and the advancement of personalized medicine. By applying the *is*SID assay, we have uncovered previously inaccessible data, providing novel insights into pathological mechanisms underlying neurodegeneration and offering potential applications for the development of diagnostic tools. Successfully applied to *in vitro* models, this approach offers further potential to elucidate seeding mechanisms at the subcellular level. For example, in combination with electron microscopy, this assay could identify the organelles and cellular structures involved in protein aggregation. Similarly, the *is*SID assay may serve as a valuable tool for studying maturation of pathology in animal models, enabling investigations into whether punctate seeding foci evolve into mature α-syn inclusions. Moreover, *in vitro* systems could be leveraged to evaluate the efficacy of therapeutic compounds targeting pathological seeding. A similar rationale can be applied at the subcellular level – if seeding activity is shown to be predominately associated with specific organelles, targeted therapeutic strategies could selectively disrupt those. Additionally, the integration of immunodetection and his-tagged monomers could enhance the specificity and sensitivity of RT-QuIC performed on bulk tissue, enabling the delineation of newly formed aggregates (containing the his-tag) in combination with other analytical techniques. Importantly, we have successfully shown that the novel *is*SID assay can be adapted to understand the cellular and spatial aspect of seeding in other proteinopathies, including tauopathies but could be expanded to include for example, Aβ and TDP-43, as well as mutant protein substrates. To this respect, while shaking is most likely required to enhance tau-*is*SID, it did not significantly enhance α-syn-*is*SID. Future studies aiming to adapt the *is*SID assay for other proteinopathies will likely require a flexible platform to allow the optimization of the incubation parameters. Our purpose-built prototypes facilitate this optimization by controlling humidity and allowing the assay to be performed in the plate reader for real time monitoring of ThT fluorescent signal.

The adaptation of *is*SID for use with tissue slides marks a significant advancement. FFPE tissue is widely available and often archived, allowing for retrospective analyses of well-characterized cases. While we acknowledge that the seeding capacity of homogenised frozen tissue might not be fully recapitulated in FFPE tissue, our results, particularly in distinguishing positivity or negativity, align with results using bulk seeding amplification assays. Furthermore, the assay’s applicability was validated in cellular models, ruling out the possibility that the *is*SID signal as an artefact associated with FFPE tissue. Another important consideration is the variability in fixation times and tissue processing protocols across brain banks. Prolonged fixation can lead to protein degradation or reduced extractability, potentially introducing inconsistencies in the seeding activity. Although we have not seen any correlation with post-mortem interval nor fixation times, performing this assay on frozen slides could be informative. Additionally, as the *is*SID assay uses immunodetection such as IHC, which relies on signal amplification, the size of the observed seeding foci observed may not directly reflect that occurring *in situ*. Comparisons between α-syn seeding foci detected by our novel assay and IHC could help clarify this. A detailed comparison of the *is*SID signal with the conventional RT-QuIC assay is also warranted. This could determine if the load of seeding competent lesions in *is*SID correlates with kinetic parameters of bulk RT-QuIC and their relative specificity and sensitivity, such as if single seeding competent lesion identified by *is*SID would produce a visible increase in bulk ThT fluorescence.

In conclusion, the novel *is*SID assay described herein offers a streamlined approach enabling the visualization of seeding activity with unprecedented spatial and cellular resolution. This significant advancement provides valuable insights into the molecular underpinnings of disease pathogenesis and progression in neurodegenerative disorders, facilitating a deeper understanding of the dynamics of protein aggregation and misfolding.

## Acknowledgments

We would like to thank the Multiple Sclerosis and Parkinson’s UK Tissue Bank at Imperial College London for their help and for providing the tissue samples uses in this study, as well as the donors and their families. We also wish to thank Dr. Francesco Aprile and Prof. Oscar Ces for their guidance and critical opinion during the development of the assay.

The technology contained in this publication is owned by Imperial College London and claimed by a pending priority patent application with the UK Intellectual Property Office (Application Number GB2410222.0)

## Author contributions

JAA conceived and conceptualized the *is*SID assay. JAA and MOJ developed the α-syn-*is*SID assay and conceptualized the manuscript. JAA and SJ developed the tau-*is*SID assay. JAA, MOJ, MJW and SJ designed the experimental plan, conducted the experiments and analyzed the data. JAA and DM conceived and conceptualized the set up (roofed chambers, plate holders and cylinders) for the *is*SID assay. DM fabricated the set up. SGR contributed to the experiments and data analysis. GG and LAH assisted with the confocal imaging. TV and BS assisted with microscopy experiments and manuscript revision. DG helped with the selection of the incidental cases. SG assisted with the experiment interpretation and manuscript writing. JAA supervised the study. JAA, MOJ, MJW, SJ, SGR and SG wrote the manuscript. All authors read and approved the final manuscript.

