## Supplementary Information for "Novel in situ seeding immunodetection assay uncovers neuronal-driven alpha-synuclein seeding in Parkinson’s disease"

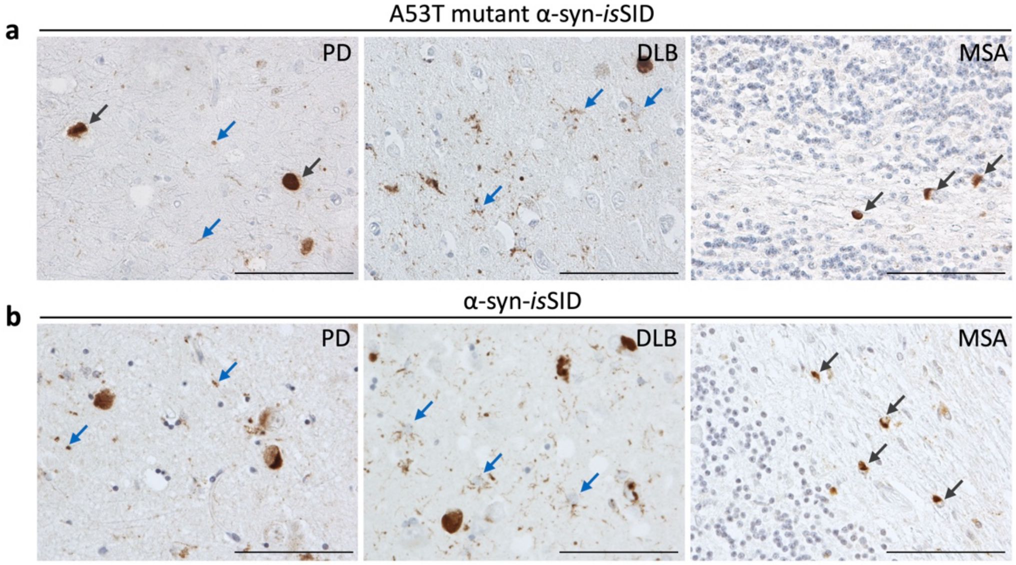


**Supplementary Fig. 1: α-Syn-*is*SID using A53T mutant α-syn protein in PD, DLB, and MSA cases**. LB (grey arrow) and dot-like/neuritic pathology (blue arrow) were seen in PD. In DLB cases, astrocytic α-syn pathology is depicted (blue arrows). GCIs were detected in MSA (grey arrows) (a). A comparison of results with α-syn-*is*SID is shown (b). Scale bar is 100 µm.

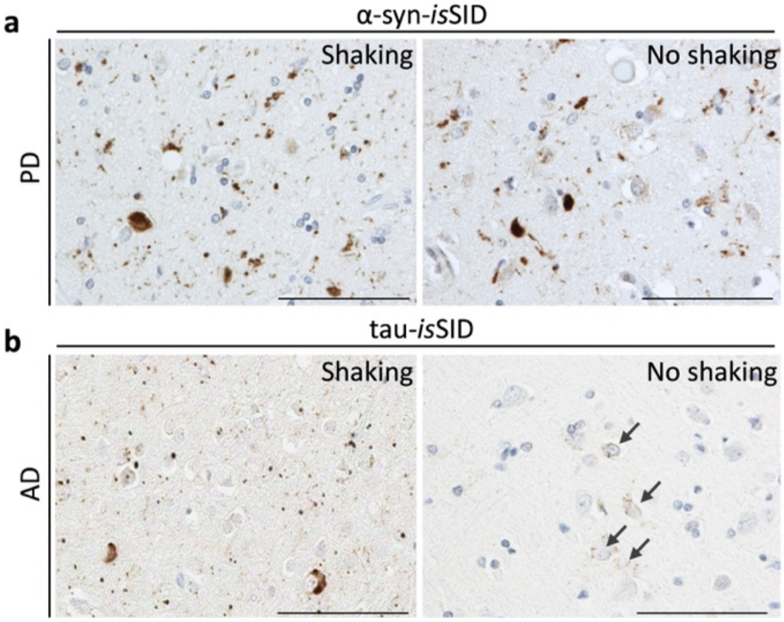


**Supplementary Fig. 2: Representative images with shaking on α-syn-*is*SID and tau-*is*SID assays.** Shaking had no impact on α-syn-*is*SID signal (**a**). In contrast, tau-*is*SID signal was lower under non-shaking conditions, although dot-like neuropil and punctate cytoplasmic staining remained detectable (**b**). Scale bar is 100 µm.

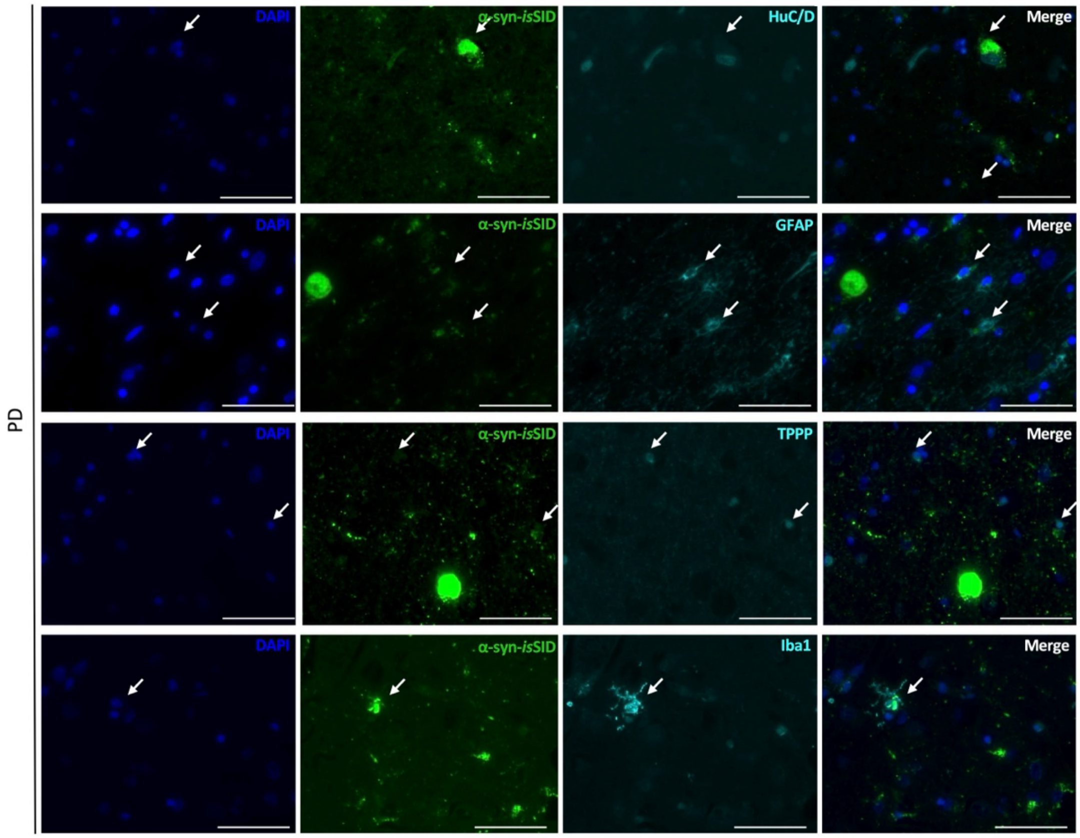


**<u>Supplementary Fig. 3</u>: Immunofluorescence confirmed seeding capacity of neuronal, astrocytic, oligodendrocytic, and microglial α-syn inclusions.** Neuronal, astrocytic, oligodendrocytic, and microglial co-localization of α-syn-*is*SID signal was confirmed using HuC/D, GFAP, TPP, and Iba1 markers, respectively. DAPI was used as a nuclear marker. Arrows depict areas of co-localization Scale bar is 100 μm.

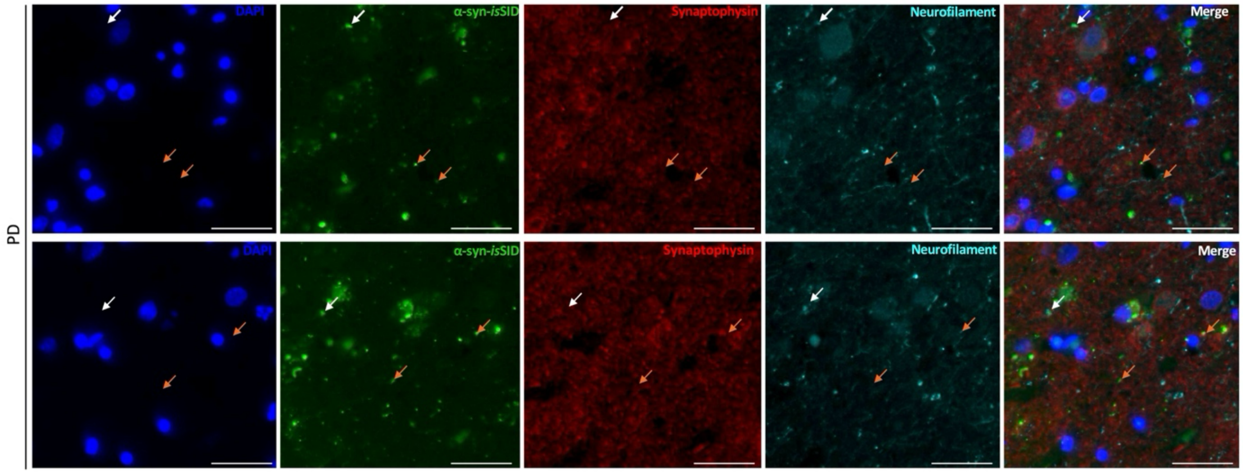


**<u>Supplementary Fig. 4</u>: Immunofluorescence shows dot-like pathology in the synapses and in close proximity.** Dot-like α-syn-*is*SID signal co-localized with pre-synaptic markers (synaptophysin) and neuronal marker (neurofilament heavy), being observed proximal to the synapses. Arrows depict areas of co-localization with synaptophysin (orange) and neurofilament heavy (white). Scale bar is 100 μm

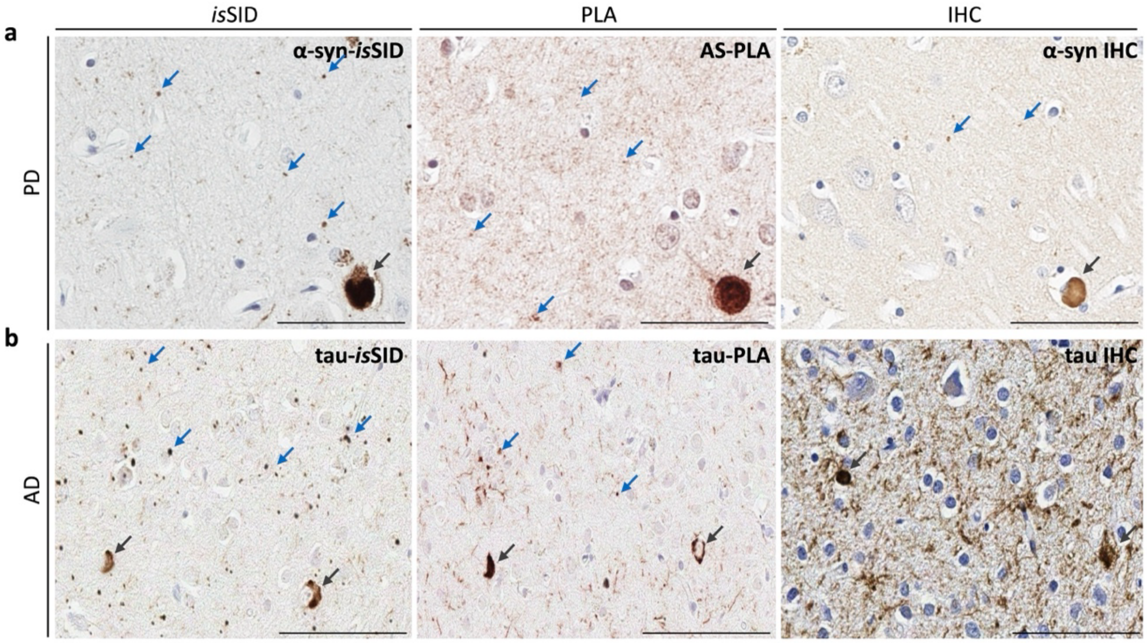


**<u>Supplementary Fig. 5</u>: Comparison of signal detected by α-syn or tau isSID, PLA, and IHC**. α-Syn aggregates, such as LBs, were detected by α-syn-isSID, AS-PLA, and α-syn IHC. However, α-syn dot-like pathology revealed by α-syn-isSID was more readily detected by AS-PLA than α-syn IHC in PD cases (a). Similar, tau aggregates were detected by all tau-isSID, tau-PLA and tau IHC, although tau dot-like pathology revealed by tau-isSID coincided with tau-PLA signal (b). Scale bar is 100 μm.
